# Dynamic autonomic nervous system patterns differentiate human emotions and manifest in resting physiology

**DOI:** 10.1101/2021.06.14.448456

**Authors:** Lorenzo Pasquini, Fatemeh Noohi, Christina R. Veziris, Eena L. Kosik, Sarah R. Holley, Alex Lee, Jesse A. Brown, Ashlin R. K. Roy, Tiffany E. Chow, Isabel Allen, Howard J. Rosen, Joel H. Kramer, Bruce L. Miller, Manish Saggar, William W. Seeley, Virginia E. Sturm

## Abstract

Whether activity in the autonomic nervous system differs during distinct emotions remains controversial. We obtained continuous multichannel recordings of autonomic nervous system activity in healthy adults during a video-based emotional reactivity task. Dimensionality reduction revealed five principal components in the autonomic time series data, and these modes of covariation differentiated periods of baseline from those of video-viewing. Unsupervised clustering of the principal component time series data uncovered separable autonomic states that distinguished among the five emotion-inducing trials. These autonomic states were also detected in baseline physiology but were intermittent and of smaller magnitude. Our results suggest the autonomic nervous system assembles dynamic activity patterns during emotions that are similar across people and are present even during undirected moments of rest.

**One Sentence Summary:** Dynamic autonomic patterns distinguish among emotions and are evident in resting physiology.

## Main Text

For over a century, one of the most contentious areas in emotion research has been the role of the autonomic nervous system (ANS). Some theorists maintain the ANS lacks precision, turning on and off during emotions in an all-or-nothing fashion *(1*), but others assert its influence is more refined, orchestrating unique visceral and motor cascades across the body during each emotion *(2, 3)*. The ANS is equipped to produce targeted changes in specific organs and muscles through organized sympathetic and parasympathetic nervous system pathways *(2)*. Previous attempts to uncover ANS signatures of emotions, however, have yielded inconclusive evidence *(4, 5)*. Some laboratory-based studies of emotional reactivity have found dissociable ANS patterns for certain negative and positive emotions *(6–9)*. Large meta-analyses, in contrast, have failed to find a robust ANS signature that is unique to each emotion *(5, 10)*. Substantial methodological differences across studies (e.g., which emotions are elicited and of what intensity, which ANS measures are collected and over what time period) *(11)* may obfuscate clear distinctions among emotions. Nonetheless, emotion-specific ANS patterns within a study—as well as across studies—remain elusive.

Here, we provide evidence that emotions are accompanied by patterned cascades of ANS activity that unfold over time *(2, 12, 13)*. Most prior studies searched for biological distinctions among emotions by collapsing dynamic ANS data into static “snapshots” that reflect average activity during various conditions *(4, 5)*. Quantifying emotional reactivity as a change score (mean activity during an emotion-inducing task minus mean activity during a baseline period), although straightforward and efficient, reduces complex physiological outflow into a single value and ignores the rich temporal structure of the ANS signals. From the spontaneous firing rates of neurons *(14)* to the intrinsic connectivity of distributed brain networks *(15)*, biological systems have dynamic properties that are critical for producing complex behaviors. We reasoned that the ANS would be no different *(16)* and that its moment-to-moment fluctuations would be essential to consider when searching for physiological patterns during emotions.

To test this hypothesis, we obtained continuous multichannel recordings of ANS physiology in 45 well-screened healthy adult volunteers (table S1) during a laboratory-based emotional reactivity task (Fig. 1A). Participants completed five “emotion trials” in which they viewed emotionally evocative videos (selected to elicit awe, sadness, amusement, disgust, and nurturant love), each approximately 90 seconds in length. Each video was preceded by a 61-second pre-trial baseline period and followed by a 31-second post-trial baseline period (herein, referred to together as the “baseline periods”) during which participants viewed an “X” on the computer monitor and were asked to clear their minds (Fig. 1B). This experimental sequence allowed participants to alternate between periods of resting fixation and periods of positive and negative emotions. At the end of each post-trial baseline, participants answered questions about what they saw and how they felt while viewing the video. All participants attended to the videos (table S2) and reported experiencing the target emotions (table S3). Whereas participants tended to report one type of emotional experience (i.e., disgust or sadness) for the negative emotion trials, they often endorsed the target emotion (i.e., awe, amusement, or affection) as well as other types of positive emotional experience for the positive emotion trials, consistent with prior studies *(6)*.

**Fig. 1.**
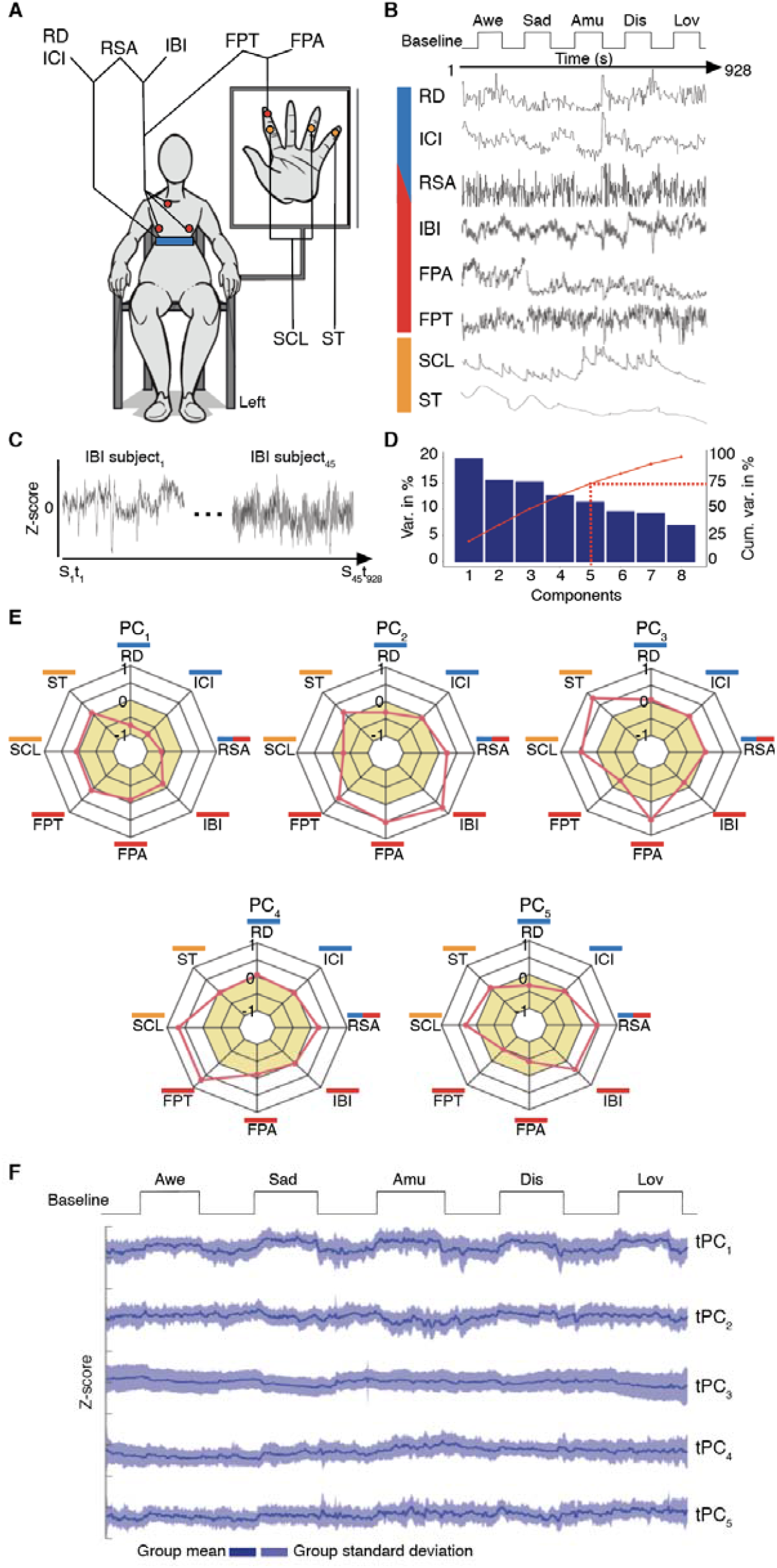
Principal components of ANS activity during the emotional reactivity task. **(A)** Multichannel recordings of ANS activity were obtained in each participant: inter-beat interval (IBI), respiratory sinus arrhythmia (RSA), finger pulse amplitude (FPA), finger pulse transit time (FPT), inter-cycle interval (ICI), respiration depth (RD), skin conductance level (SCL), and skin temperature (ST). Blue indicates respiratory; red, cardiovascular; and orange, dermal signals. **(B)** Continuous recording of ANS activity were obtained throughout the emotional reactivity task, during which participants viewed five emotionally evocative videos (awe [Awe], sadness [Sad], amusement [Amu], disgust [Dis], and nurturant love [Lov]). Each video was preceded by a pre-trial baseline and followed by a post-trial baseline (Baseline) during which they viewed an “X” on the computer monitor. **(C)** The ANS time series data were *z*-scored in each participant and then concatenated across individuals. **(D)** A principal component analysis revealed five principal components (PCs), which explained 75% of the variance in the ANS time series data during the emotional reactivity task. **(E)** Radial plots for PC_1-5_ show each PC’s eigenvector loadings. For positive eigenvector loadings, higher values reflect greater IBI (slower heart rate), FPA (larger pulse amplitude in the finger), FPT (slower transmission of pulse from heart to finger), SCL (higher electrodermal activity), ST (higher skin temperature on the finger), RD (greater respiration depth), ICI (slower respiration rate), and RSA (higher heart rate variability). For negative eigenvector loadings, the opposite patterns were true. Radians in ochre represent negative loadings. **(F)** The time series of each PC (tPC) were computed and plotted to illustrate second-by-second fluctuations during the emotional reactivity task.

Throughout the emotional reactivity task, we measured eight channels of ANS activity: inter-cycle interval (a measure of respiration rate), respiration depth, inter-beat interval (a measure of heart rate), finger pulse amplitude, finger pulse transit time, respiratory sinus arrhythmia, skin conductance level, and skin temperature. This broad array of respiratory, cardiovascular, and electrodermal measures enabled us to capture a wide range of ANS activity representing both sympathetic and parasympathetic nervous system influences *(17)*. After standard preprocessing of the ANS time series, second-by-second averages for each channel were exported, and missing values were spline interpolated (depending on the channel, between 0.0-5.5% of total data, see fig. S1). To transform the ANS measures into a standard scale, we converted the second-by-second averages to *z*-scores within each participant and then concatenated the time series data for each channel across the participants (Fig. 1C).

We used principal component analysis *(18)*, a dimensionality reduction technique that uncovers latent modes of covariation, and identified five principal components (PCs) in the concatenated ANS time series data. Each PC explained at least 10% of the total variance, and, together, they explained 75% (see Fig. 1D and fig. S2). The PCs lacked one-to-one mappings with the cardiovascular, respiratory, and dermal signals, which suggested a more complex ANS organization during the emotional reactivity task. Activity in some channels, however, loaded more strongly on certain PCs than on others: respiratory activity (i.e., respiration depth and inter-cycle interval) loaded strongly on PC_1_; cardiovascular activity (i.e., finger pulse amplitude and inter-beat interval), on PC_2_; dermal activity (i.e., skin temperature and skin conductance level), on PC_3_; cardiovascular and dermal activity (i.e., finger pulse transit time and skin conductance level), on PC_4_; and inter-beat interval and skin conductance level, on PC_5_ (Fig. 1E).

To investigate this latent structure in more detail, we next computed the time series of each PC (tPC). A tPC was calculated by multiplying the standardized second-by-second data in each ANS channel by its PC loadings and then summing the weighted time series of the individual channels. We repeated this approach for each of the PCs, which yielded five tPCs with distinct temporal dynamics (Fig. 1F). We then reduced the tPCs during the baseline periods and each emotion trial into static averages and compared their overall magnitudes. Analyses of variance and multinomial logistic regression analyses confirmed that the mean tPC amplitudes differed among the emotion trials (fig. S3A-G). Consistent with prior studies, these results suggested there were some reliable differences in mean ANS activity among distinct emotions (fig. S3) *(4, 6, 8)*. Although our models reached above-chance classification probabilities, we expected more robust ANS differences among the emotion trials may be embedded within the temporal dynamics of the tPCs.

In our next analyses, we preserved the second-by-second fluctuations of the tPCs to investigate whether they offered additional insights into the architecture of ANS patterning. Examination of the group-averaged tPCs suggested that activity in tPC_1_ aligned with the task structure and showed comparable increases during all the emotion trials (Fig. 2A). The magnitude of tPC_1_ was more negative during the baseline periods (i.e., reflecting slower, deeper respiration and slower heart rate) and more positive during the emotion trial (i.e., reflecting faster, shallower respiration and faster heart rate; fig. S4). The other tPCs, however, did not show a similar time course but instead exhibited more complex fluctuations, especially during the emotion trials (fig. S5).

**Fig. 2.**
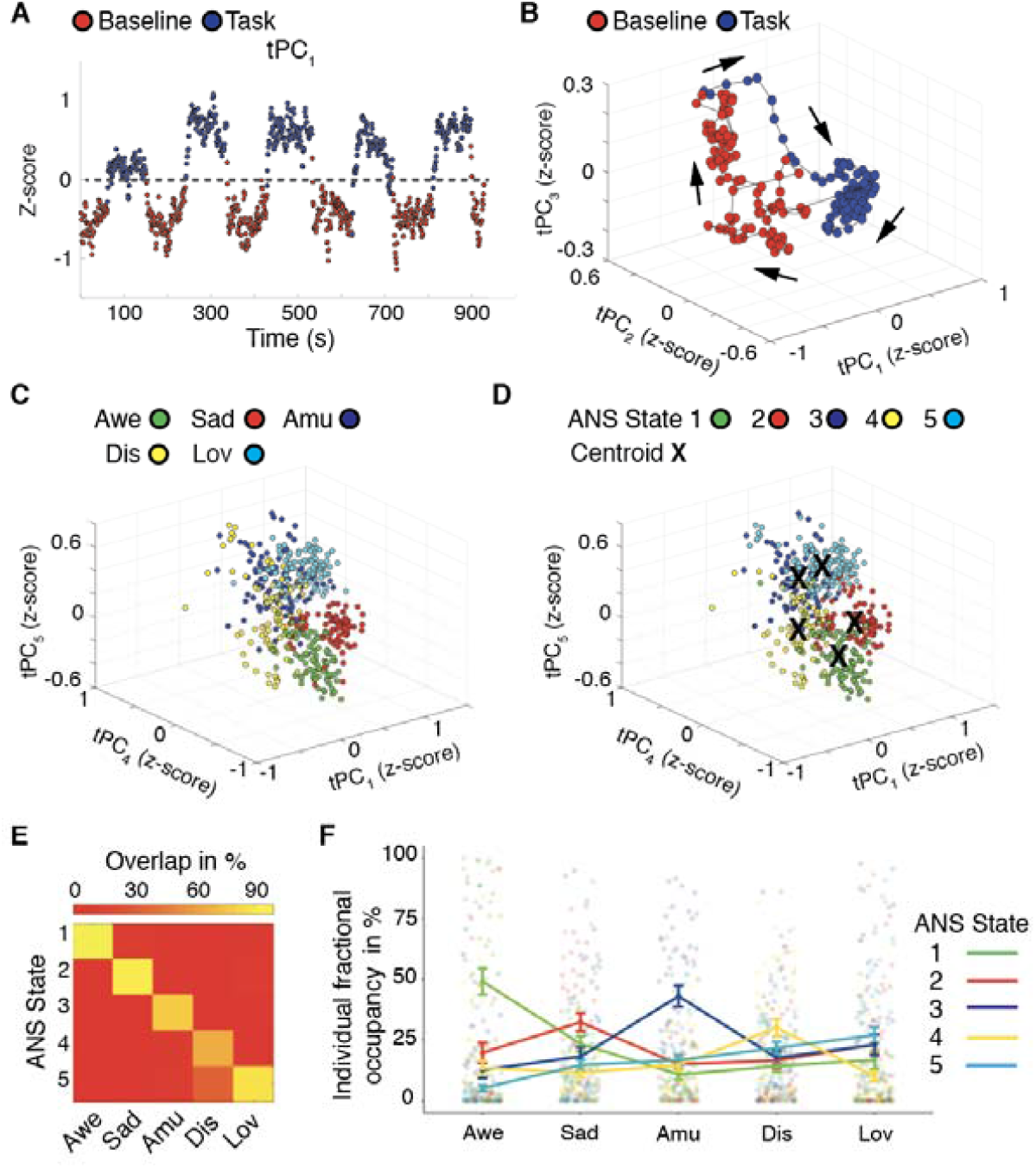
Dynamic ANS states emerged from continuous physiological measures. **(A)** During the emotional reactivity task, the amplitude of the time series of principal component 1 (tPC_1_) fluctuated over time and differentiated baseline periods (“Baseline,” shown in red) from the emotion trials (“Task,” shown in blue). **(B)** When projected into a low-dimensional manifold, the flow of the tPCs within the embedding space (depicted by arrows) separated the baselines from the emotion trials. Periods of the tPCs overlapping only with the emotion trials were then plotted in the low-dimensional space. Color-coding each data point by **(C)** the emotion trial in which it was acquired or **(D)** its assigned ANS state following k-means clustering suggested a remarkable similarity between the spatial topography of these two plots. **(E)** A confusion matrix revealed that during each emotion trial, there was a single predominant ANS state (except for the disgust trial, which had two predominant ANS states), as indicated by the percentage of tPC data point during each trial that aligned with each ANS state. **(F)** To confirm that the group-level results were consistent with patterns found in individuals, we assigned data points in the tPCs of th individual participants to the closest cluster and computed fractional occupancy scores for each ANS state in each emotion trial, as shown in the line-plots with associated standard error bars. As in the group-level analysis, individual participants tended to occupy a predominant ANS state during each emotion trial, and the state that they occupied in a given trial was similar across individuals.

We modeled the temporal trajectories of the five tPCs during the baseline periods and the emotion trials in a low-dimensional manifold, a representation that can further elucidate underlying structures in multidimensional data *(19, 20)*. As above, the overall trajectory of ANS activity in the low-dimensional manifold separated the baseline periods from the emotion trials (Fig. 2B), which confirmed physiological responsivity during video-viewing *(19, 20)*. Consistent with theories that emphasize arousal as a central dimension of emotions *(5, 10)*, some ANS changes (e.g., respiration depth) differentiated the emotion trials from the baseline periods but did not distinguish among the emotion trials themselves.

Next, we removed the baseline periods to inspect the temporal dynamics of the tPCs during the emotion trials alone, a more rigorous search for ANS patterning. We plotted the tPCs in the low-dimensional space and labeled each data point by the emotion trial in which it was acquired. Remarkably, this plot revealed five dynamic ANS patterns, each aligning with a different emotion trial (Fig. 2C). We next applied unsupervised k-means clustering, an approach agnostic to the temporal order of the data *(21)*, to the tPCs to confirm the presence of distinct ANS patterns during the emotion trials. This technique also uncovered five clusters or “ANS states” in the tPCs, a solution that was further supported by a silhouette analysis (fig. S6). When plotted in the low-dimensional space (Fig. 2D), the spatial topography of these ANS states largely mirrored that which emerged when the tPC data points were instead labeled with their attendant emotion trial. Indeed, each ANS state had a predilection for a specific emotion trial (Fig. 2E). ANS State 1 arose exclusively during the awe trial (100%); ANS State 2 arose most often during the sadness trial (92%); ANS State 3, during the amusement trial (78%); ANS State 4, during the disgust trial (67%); and ANS State 5, during the disgust (33%) and nurturant love (87%) trials.

As the ANS states were derived from the time series data averaged across the sample, we next examined the degree to which these states manifested in individual participants. We identified the cluster-centroid of each ANS state, which is considered its prototype, and assigned every data point in the tPCs of the individual participants to the closest state based on Euclidean distance. We calculated fractional occupancy scores, the percentage of time spent in each ANS state during each emotion trial, for every participant. A two-way analysis of variance comparing the fractional occupancy scores across the emotion trials found a main effect of ANS state, *F*(4,1100)=5.1, *p*<0.0005, and an interaction between ANS state and emotion trial, *F*(16,1100)=13.2, *p*<0.0005. This analysis indicated that, as found at the group level, individual participants most often occupied ANS State 1 during the awe trial, ANS State 2 during the sadness trial, ANS State 3 during the amusement trial, ANS State 4 during the disgust trial, and, to a lesser extent, ANS State 5 during the nurturant love trial (Fig. 2F, *p*<0.05 Bonferroni-corrected *t*-tests). As an additional test, we conducted k-means clustering on the tPCs; here we limited each analysis to an individual participant’s data. By computing the Euclidean distance between the cluster-centroids in each participant and those derived at the group level, we again assigned each second of the tPCs to the corresponding ANS state. Like the results conducted across the sample, these analyses indicated that a single ANS state, identifiable by its unique physiological profile (fig. S7), was predominant during each emotion trial (fig. S8).

Functionalist theories propose emotions are accompanied by patterned suites of ANS and motor changes that evolved to address recurrent problems in life *(22)*. Consistent with this view, our results suggested dynamic ANS patterns differentiated among the emotion trials. By disrupting homeostasis, emotions are thought to organize activity across bodily systems (within the ANS and between the ANS and other systems) that are typically uncoordinated at rest *(2, 22, 23)*. Although maintenance of regular physiological rhythms is a mainstay of homeostasis, even at rest there is unexplained variability in ANS outflow that adds complexity to an otherwise steady state *(24, 25)*. Thus, we wondered whether the ANS patterns that emerged during the emotion trials might be detectable within the cacophony of undirected physiology. To investigate this possibility, we performed a principal component analysis on the standardized ANS time series data, concatenated across the sample, from a two-minute resting period that preceded the emotional reactivity task. This analysis found PCs with loadings that were similar to those found during the emotional reactivity task (Fig. 3A).

**Fig. 3.**
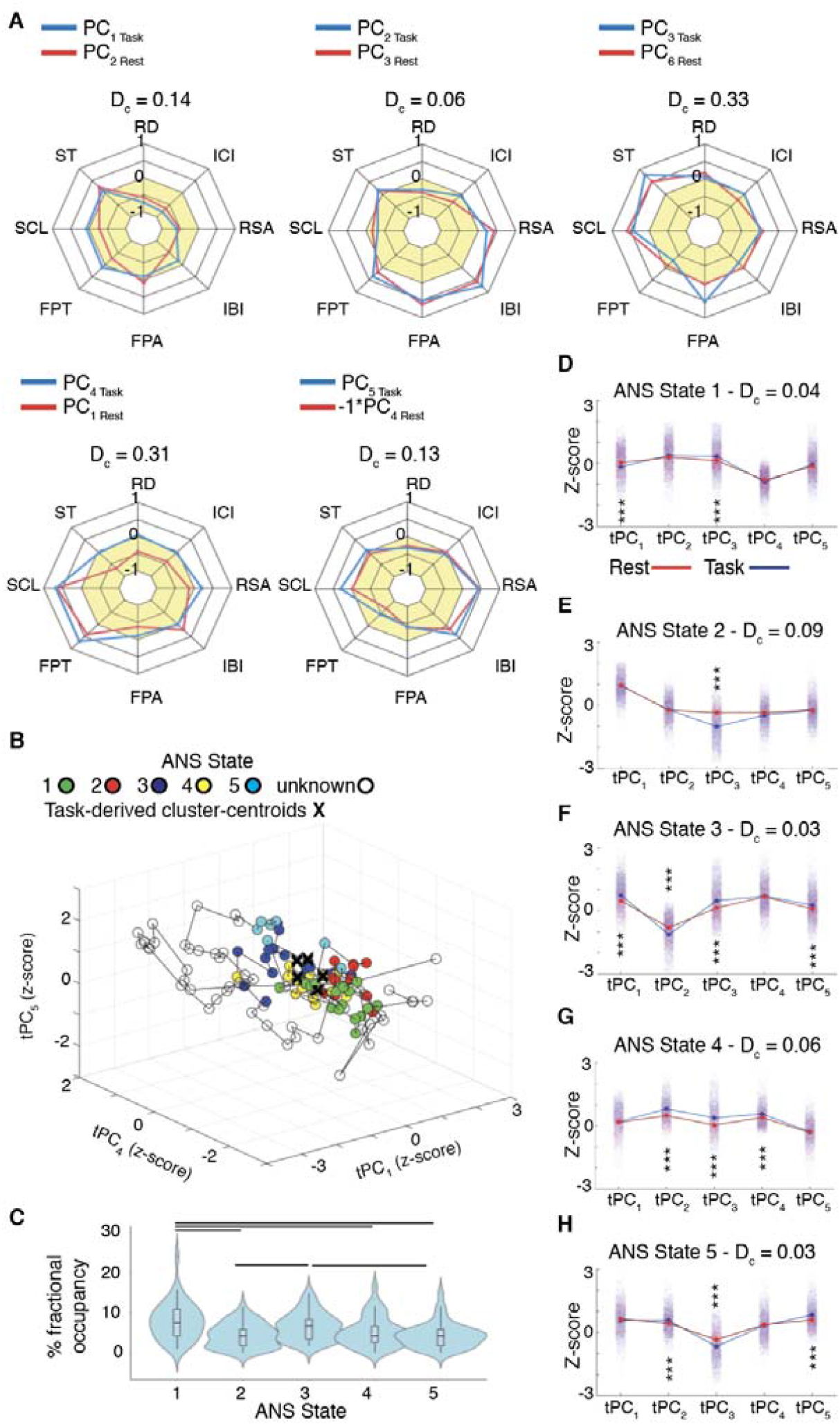
Emotion-relevant ANS states were detectable in resting physiology. **(A)** We performed a principal component analysis on the ANS recordings acquired during the two-minute resting period. Measures of cosine distance (D_c_) confirmed the principal components (PCs) from the resting period (“Rest,” red lines) resembled those from the emotional reactivity task (“Task,” blue lines). **(B)** To illustrate the spatial topography of resting ANS activity in the low-dimensional space, the projected time series of the principal components (tPCs) are shown for a representative participant. Data points from the resting period that were assigned to one of the five ANS states from the emotional reactivity task are color-coded as in Fig. 2D; data points greater than one standard deviation from the mean distance of a cluster-centroid were classified as unknown states. **(C)** Fractional occupancy scores reflect the percentage of the resting period that participants spent in ANS states that aligned with those from the emotional reactivity task. During the resting period, participants spent the greatest percentage of time in ANS States 1 and 3, states that were predominant during the awe and amusement trials. Bars reflect *p*<0.05, Bonferroni-corrected *t*-tests. **(D-H)** Line-plots, with jitter depicting the averaged magnitude of the tPCs, show a high degree of similarity between the mean tPC activity during the ANS states during the resting period (red lines) and the emotional reactivity task (blue lines), as measured by D_c_; *** *p*<0.05, Bonferroni-corrected *t*-tests.

As before, we generated the corresponding tPCs and computed the Euclidean distance from each data point to the cluster-centroids from the emotion trials. Here, these distances reflected the similarity between each second of the resting period tPCs and the emotion-relevant ANS states. We assigned data points near a cluster-centroid to the closest ANS state; data points that did not align with one of the cluster-centroids remained unassigned (Fig. 3B, fig. S10 and S11). During the resting period, participants spent 7-57% in an ANS state that resembled those from the emotional reactivity task (Fig. 3C and fig. S10). A one-way analysis of variance, *F*(4,220)=8.1, *p*<0.05, with Bonferroni-corrected pairwise *t*-tests *(p*<0.05) found that, on average, participants spent more time in ANS State 1 (7.8%) and ANS State 3 (6.5%) than in ANS State 2 (4.3%), ANS State 4 (5.1%), or ANS State 5 (4.2%; Fig. 3C). When considering each ANS state’s associated emotion trial, these findings suggested that, at rest, participants spent more time in ANS states emblematic of awe and amusement than in those of sadness, disgust, or nurturant love. Whereas during the emotional reactivity trial the presence of the ANS states tended to be sustained, their occurrence during the resting period was brief and intermittent (Fig. 3B, fig. S10 and S11).

The presence of ANS states at rest that resembled those from the emotional reactivity task suggested an organization to basal physiological outflow that has previously gone unrecognized. We conducted additional tests to evaluate the degree of similarity between the ANS states uncovered at rest and those found during the emotion trials. Measures of cosine distance (D_c_), in which lower values indicate higher similarity, confirmed that the ANS states derived from the resting period had tPC profiles that were similar to those from the emotion trials (D_c_ State 1 = 0.04, State 2 = 0.09, State 3= 0.03, State 4 = 0.06, State 5 = 0.03, Fig. 3D-H). A three-way analysis of variance with Bonferroni-corrected pairwise *t*-tests *(p*<0.05) comparing the tPC magnitudes of the ANS states from the resting period with those from the emotion trials also found a main effect of condition (resting period versus emotion trials), *F*(1,112875)=2.6, *p*<0.05 (Fig. 3D-H). Taken together, these results indicated that the ANS states from the resting period had a similar composition to those from the emotion trials but, on average, were of a lower magnitude.

Prior studies that have searched for ANS signatures of emotions have yielded inconsistent results. By taking a novel approach to the dimensionality reduction and analysis of multichannel ANS time series data, we found evidence for dynamic ANS patterns that differentiated among emotions and were similar across people. While different constellations of ANS activity distinguished among the trials in the emotional reactivity task, these patterns likely represent one possible manifestation—rather than the only manifestation—of each emotion. Although it seems improbable that the present study has revealed the singular physiological signature of each of these emotions, our results do suggest there are predictable ANS cascades that unfold in a similar fashion across individuals who are viewing the same evocative stimuli. This type of patterned reaction is consistent with functionalist models that propose emotions are time-tested states with unique physiological profiles in the body and brain.

The ANS, together with other systems, not only activates during salient moments but also creates the physiological backdrop on which emotions unfold. Always present, the internal milieu of the body forms the basis of ongoing feeling states (e.g., often referred to as “mood” or “core affect”) that influence cognition and shape behavior *(28)*. Our results suggest there is an intrinsic organization to resting physiology that has previously gone undetected, with patterned constellations of ANS activity arising not only during emotions but also during periods of relative quiescence. Whether these transient ANS states reflect “flickers” of emotions that color short-lived, or even more enduring, subjective experiences or reflect a fundamental “readiness” property of the ANS that prepares the organism to react to salient stimuli in a predictable way is unknown but warrants future research. Future studies that further delineate the dynamic architecture of ANS patterns—during emotions and at rest—will be critical for advancing current models of emotions neurobiology.

## Acknowledgments

We thank the participants for their contributions to research. We thank Edoardo Pasquini for his contribution as a scientific illustrator.

## Author contributions

LP: Conceptualization, Analysis, Writing. CRV, ELK, SRH, AL, JAB, ARKR, TEC, and IA: Analysis, Writing. HJR, JHK, and BLM: Conceptualization, Writing. MS, WWS, and VES: Conceptualization, Analysis, Writing.

## Data and materials availability

Data, code, and materials used in the analyses are publicly available at https://github.com/lollopasquini/Dynamic_ANS.

## Supplementary Materials

Materials and Methods

Figures S1-S11

Tables S1-S3

External Databases available at https://github.com/lollopasquini/Dynamic_ANS

References *(33-37)*

