## Supplement for "Dynamic autonomic nervous system patterns differentiate human emotions and manifest in resting physiology"

**This PDF file includes:**

Materials and Methods

Supplementary Materials

Supplementary Results

Figs. S1 to S11

Tables S1 to S3

**Other Supplementary Materials for this manuscript include the following:**

Data, code, and materials used in the analyses are publicly available at <https://github.com/lollopasquini/Dynamic_ANS>.

Materials and Methods

Participants

Fifty-nine healthy adults were recruited from the Hillblom Aging Network, a longitudinal cohort followed by the University of California, San Francisco (UCSF) Memory and Aging Center. Participants underwent a comprehensive multidisciplinary neurological, neuropsychological, functional, and neuroimaging assessment and were cognitively normal and free of current or previous neurological or psychiatric conditions. All participants had a Clinical Dementia Rating Scale score of 0 (range 0 – 3), which indicates intact daily functioning (*33*), and a Mini-Mental State Examination score of 28 or higher (range 0 – 30), which indicates intact mental status (*34*). At the time of the emotional reactivity trial, no participants were taking medications that might affect ANS physiology (i.e., stimulants, acetylcholinesterase inhibitors, or beta blockers). Race and ethnicity were self-reported by study participants, and categories were defined by investigators based on the US Office of Management and Budget’s Revisions to the Standards for the Classification of Federal Data on Race and Ethnicity. Handedness and years of education were also reported. A subset of participants was excluded from the data analyses due to incomplete physiological measures (see below), which resulted in a final sample of 45 participants. The demographic and cognitive data for the final sample are shown in supplementary table S1. Participants provided informed consent prior to participation, and the study was approved by the UCSF Human Research Protection Program.

Laboratory-Based Assessment of Emotions

*Procedure*

The laboratory-based assessment of emotions was completed at the UCSF Center for Psychophysiology and Behavior. Participants were seated in a comfortable chair in a well-lit experiment room. Sensors were applied to obtain continuous measures of physiological activity (Fig. 1A). Participants were recorded with a semi-obscured remotely controlled video camera throughout the testing session; behavior from the videos was not analyzed in the present study. Participants completed a battery of tasks designed to assess emotional reactivity, empathy, emotion regulation, and resting ANS physiology. Only data from the emotional reactivity task and resting period were included in our analyses.

*Tasks*

Emotional Reactivity Task

Participants viewed five videos (88 – 104 seconds in length) selected to elicit awe, sadness, amusement, disgust, or nurturant love. Each video was preceded by a 61-second pre-trial baseline in which they were asked to try to clear their mind and followed by a 31-second post-trial baseline in which participants viewed a black “X” on a white screen (Fig. 1B). Piloting in an independent sample indicated that the videos elicited the target emotions. After viewing each video, participants answered questions about the video and their emotional experience. All questions were presented visually on the computer monitor and audially over speakers.

Resting Period

Prior to the emotional reactivity task, participants sat quietly through a two-minute resting period in which they watched a black “X” on a white screen. They were instructed to try to clear their mind during the trial.

*Measures*

Physiological Recordings

Continuous measures of physiological activity were obtained with Biopac MP150 bioamplifiers and a computer equipped with AcqKnowledge software (v4.4, <https://www.biopac.com/>) (*17*): *(I)* Inter-beat interval (IBI): Electrodes were placed in a bipolar configuration on opposite sides of the participant's chest; the heart rate was calculated as the number of R waves from the electrocardiogram per minute. *(II)* Inter-cycle breath interval (ICI): A pneumatic bellows-based respiration transducer was stretched around the thoracic region, and the respiration rate was measured as the number of inspirations per minute. *(III)* *Respiration depth (RD)*: The point of the maximum inspiration minus the point of maximum expiration was determined from respiratory tracing. *(IV)* *Finger pulse amplitude (FPA):* A photoplethysmograph recorded the amplitude of blood volume in the finger using a photocell taped to the distal phalanx of the index finger of the nondominant hand. *(V)* *Finger pulse transit time (FPT)*: The time interval in milliseconds was measured between the *R* wave of the electrocardiogram and the upstroke of the peripheral pulse at the finger site, recorded from the distal phalanx of the index finger of the nondominant hand. *(VI)* *Skin temperature* *(ST)*: A thermistor attached to the distal phalanx of the little finger of the nondominant hand recorded temperature in degrees Fahrenheit. *(VII)* Skin conductance level (SCL): SCL, a measure of sympathetic activity (*35*), was measured through a constant-voltage device passing a small voltage between Ag/AgCl Silver 8 mm electrodes (using an electrolyte of sodium chloride) attached to the palmar surface of the middle phalanges of the ring and index fingers of the non-dominant hand. *(VIII)* *Respiratory sinus arrhythmia (RSA)*: RSA, a measure of parasympathetic activity (*36*), was calculated with the peak-valley approach as the difference in msec between the shortest inter-beat interval during inspiration and the longest inter-beat interval during expiration.

Physiological data were processed using a custom pipeline scripted in AcqKnowledge. Briefly, algorithms identified and marked the signature components of each waveform, and these marks were then visually inspected for errors and noise. Outliers in the data were considered ± 3 standard deviations from the mean level during the trial; these periods were interpolated if their duration was three seconds or less and deleted if their duration was greater than three seconds. Second-by-second averages for each channel were then exported for use in subsequent analyses.

Self-report measures

After viewing each video, participants were asked, “What happened in this movie?” and then selected their answer from multiple choice. On a three-point scale (0=*none*, 1=*a little*, 2=*a lot*), they additionally reported their experience of anger, sadness, disgust, fear, awe or amazement, love or affection, amusement or happiness, excitation or enthusiasm, embarrassment, proudness, and surprise while watching each video.

*Analyses*

R (<https://www.r-project.org/>) and MATLAB2020a (<https://www.mathworks.com/products/matlab.html>) were used for the analyses.

Emotional Reactivity Task

*Validity checks*

Correct responses to the questions about each video’s content were scored as 1, and incorrect responses were scored as 0. We used these scores to confirm that all participants had paid attention during the task (100% accuracy). For the emotional experience questions, we conducted analyses of variance to determine whether there was a main effect of emotion trial on each type of emotional experience.

*Principal component analysis*

Participants with more than 20% missing data in any physiological channel across the emotional reactivity task, as determined with the *vim* package, were excluded from our analyses; 14 out of 59 participants were excluded based on this criterion, leaving 45 participants in our final sample.

Incidental missing values were replaced through spline interpolation using the *imputeTS* package in R. Between 0.2% (IBI) and 5.5% (RSA) of missing data across the whole sample was replaced through the interpolation procedure (fig. S1). The time series of the ANS channels were then *z*-scored within each participant using the *scale* function in R (Fig. 1C), which ensured that individual differences in ANS dynamics were comparable across participants and measures. For each ANS channel, the standardized time series were then concatenated across the sample (Fig. 1C) for use in a principal component analysis using the *factoextra* package in R. The first five principal components (PCs) were selected for further analysis. Each PC was characterized by a set of eigenvector loadings that reflected the contribution of the ANS signals to each PC. The time series of each PC (tPC), which were comprised of the second-by-second dot products of the eigenvector scores and the standardized ANS signal, were also computed. Each tPC reflected the continuous fluctuations in each PC’s amplitude across the emotional reactivity task.

*Analysis of variance*

For each participant, we computed the average magnitude of each tPC during the baseline periods and each emotion trial. We conducted five analyses of variance (one for each tPC) with post hoc Bonferroni-corrected *t*-tests (*p*<0.05) to assess whether the average tPC magnitudes differed across the emotion trials (*aov* and *t.test* packages in R).

*Multinomial logistic regression*

To assess whether each emotion trial was characterized by a distinct pattern of tPC activity, we performed multinomial regression analyses (*37*) using the R packages *nnet* and *caret*. In model_1,_ we tested whether the average tPCs from each emotion trial predicted the trial in which they were acquired (i.e., baseline periods averaged across trials, awe, sadness, amusement, disgust, or nurturant love). In model_2_, we omitted the baseline periods and included only the data from the emotion trials, which allowed us to conduct a more stringent test of whether the mean tPCs could differentiate among the emotion trials alone. A 10-fold cross-validation approach was used to derive a confusion matrix for each model, which reflected the correct classification of each emotion trial and provided measures of sensitivity, specificity, and accuracy.

*Low-dimensional manifold*

To investigate the trajectory of the tPCs within a low-dimensional embedding space (*19*), we created a topological manifold (*20*). Manifolds facilitate the exploration of dynamic systems and have been used to uncover neural mechanisms of complex human and non-human animal behaviors (*19*, *20*). To compute representative trajectories for the tPCs, we created two separate matrices in which the columns stored the second-by-second tPC data from the baseline periods or the emotion trials. Given that the trials in the emotional reactivity task differed in length, we used linear interpolation (*fillmissing* function in MATLAB R2020a) to derive time series of the same length, matched to the duration of the longest trial. In each matrix, we averaged across the columns to compute the mean time series of each tPC across the baseline periods and the emotion trials and then plotted the trajectories into the embedding space.

*K-means clustering and ANS state analyses*

We performed k-means clustering (*kmeans* function in MATLAB R2020a) (*21*) with 10,000 iterations and 10 replications on the group-averaged tPCs from the emotion trials, which yielded clusters of dynamic ANS activity. The optimal number of clusters, or ANS states, was confirmed with a silhouette analysis. Each data point (i.e., each second of the time series data) was assigned to the nearest cluster-centroid—the arithmetic mean of all of the time points that belong to that cluster—using Euclidean distance. A cross-tabular confusion matrix was used to investigate the percentage of seconds of each emotion trial that were assigned to each ANS state.

We next examined whether the ANS states found in the tPC data averaged across the sample were also evident in individual participants. For each participant, we assigned every tPC data point during the emotion trials to the ANS state with the closest cluster-centroid, as measured by Euclidean distance. We used each participant’s resultant ANS state occupancy time series to compute the percentage of data points from each emotion trial that were assigned to each ANS state, a measure of ANS state fractional occupancy. We then conducted a two-way analysis of variance (*p*<0.05) to determine whether there were differences in ANS state fractional occupancies across the emotion trials (*aov* package in R). We also used a two-way analysis of variance with a random intercept for subject nested in the factor signal (*p*<0.05) to compare the amplitude of the physiological signals across the ANS states (*lmer* package in R).

We performed additional control analyses in which we conducted k-means clustering on the tPCs from the emotion trials in individual participants. The ANS states that were identified in individual participants were then aligned to those found at the group level based on the Euclidean distance between the individually generated cluster-centroids and the group-level cluster-centroids (*pdist* function in MATLAB R2020a). We again used a two-way analysis of variance to determine whether there were differences in ANS state fractional occupancies across the emotion trials (*aov* package in R).

Resting Period

*Principal component analysis of resting period ANS time series*

The ANS time series during the two-minute resting period were preprocessed with the same procedure as those from the emotional reactivity task. We then conducted a principal component analysis on the standardized ANS time series data from the resting period, concatenated across participants. Computing separate principal component analyses on two distinct data sets can yield similar components that have a different sign or order of eigenvector loadings. To identify correspondent PCs across the emotional reactivity task and the resting period, eigenvector loadings of individual PCs were compared across decompositions using cosine distance. Cosine distance was chosen because it is better suited than other distance measures to compare the overall patterns in magnitude change shared by two independently generated, short vectors. We reordered the PCs from the resting period analysis in such a way to maximize the similarity with the eigenvector loadings of PCs derived from the emotional reactivity task (*pdist* function in MATLAB R2020a). This also involved iteratively assessing the similarity between the eigenvector loadings from the emotional reactivity task decomposition and the inverse of the eigenvector loadings derived from the resting period decomposition. Once the most similar PCs were identified, we vectorized the eigenvector loading matrices and used Pearson’s correlation coefficients (*p*<0.05; *corr* function in MATLAB R2020a) to examine the correspondence between the PCs derived from the resting period and those from the emotional reactivity task. We then generated tPCs for the resting period by using the rearranged eigenvector loadings derived from the principal component analysis of the resting period ANS time series.

*ANS state extraction from the resting period*

To determine the degree to which the ANS states from the resting period resembled those from the emotional reactivity task, we computed the Euclidean distance between each data point in the tPCs from the resting period and the cluster-centroids of the ANS states from the emotion trials. Data points in close proximity to a cluster-centroid (less than one standard deviation from the mean distance of every resting period time point to any cluster-centroid) were assigned to the nearest ANS state. We then computed the ANS state fractional occupancy for each participant, which was the percentage of seconds in the resting period aligned with each ANS state. We used a one-way analysis of variance, with Bonferroni-corrected post hoc *t*-tests (*p*<0.05; *aov* package in R) to compare the ANS state fractional occupancies of participants. We used cosine distance to compare the mean magnitude of the tPCs in each ANS state during the resting period with those during the emotional reactivity task (*pdist* function in MATLAB R2020a). Again, cosine distance was chosen because it is ideally suited to compare the overall patterns in magnitude change shared by two short vectors. We then performed a three-way analysis of variance (*p*<0.05) to assess whether the mean magnitudes of the tPCs in each ANS state differed between the resting period and the emotional reactivity task (*aov* package in R). For each ANS state, the tPC magnitudes from the resting period and the emotional reactivity task were compared with two-sample *t*-tests (*p*<0.05, Bonferroni-corrected for a total of 25 pairwise comparisons; *t.test* package in R).

Supplementary Results

Emotional Reactivity Task

*Self-Report Measures*

Video content

All participants provided correct responses to the questions about the content of each video, which confirms they paid attention throughout the emotional reactivity task (table S2).

Emotional experience

Participants reported experiencing moderate levels of the target emotion during each video in the emotional reactivity task. During the awe trial, experience of awe was the most intense followed by amusement; during the sadness trial, experience of sadness was most intense followed by affection; during the amusement trial, experience of amusement was most intense followed by pride; during the disgust trial, experience of disgust was most intense followed by surprise; during the nurturant love trial, experience of amusement was most intense followed by affection (table S3).

*Physiological Activity*

Dynamic ANS activity characterized PC_6-8_

In our primary analyses, we retained the first five PCs from the group-averaged ANS time series data. PC_6-8_ did not explain a substantial portion of variance (<10%) (fig. S2A) and, thus, were not examined further. The eigenvector loadings of PC_6-8_ were weaker than those from PC_1-5_. Inter-beat interval, inter-cycle interval, and skin conductance level positively loaded on PC_6_; skin temperature and finger pulse amplitude negatively loaded on PC_7_; and inter-cycle interval positively and respiration depth negatively loaded on PC_8_. A visual inspection of each PC’s time series suggested that each had its own unique temporal dynamics over the course of the emotional reactivity task (fig. S2B).

Analyses of variance distinguished among the emotion trials using the mean tPC magnitudes

Five one-way analyses of variance were performed to assess whether the mean amplitudes of the tPCs differed across the emotion trials. These analyses revealed a main effect of emotion trial for the amplitudes of tPC_1_, *F*(1,268)=73.8, *p*<0.0001; tPC_4_, *F*(1,268)=17.0, *p*<0.0001; and tPC_5_, *F*(1,268)=39.0, *p*<0.0001. For tPC_3_, the main effect of emotion trial approached significance, *F*(1,268)=3.6, *p*<0.06, but the main effect was not significant for tPC_2_, *F*(1,268)=0.5, *p*=0.496. Violin plots and post hoc *t*-tests (*p*<0.05, Bonferroni-corrected for multiple comparisons) revealed that, on average, the magnitude of the tPCs differed across the emotion trials (fig. S3A-E).

Multinomial logistic regression models separated the emotion trials

The 10-fold cross-validated multinomial regression analyses for model_1_ (fig. S3F) and model_2_ (fig. S3G) revealed above-chance classification probabilities with modest to high levels of sensitivity, specificity, and accuracy (mean and confidence interval for model_1_ = 0.49 [0.45, 0.52] and model_2_ = 0.44 [0.40-0.48]). These findings offered additional support that the averaged magnitudes of the tPCs distinguished among the emotion trials.

Amplitude of physiological channels and tPC_1_ magnitude

Having determined that the magnitude of tPC_1_ varied with the structure of the emotional reactivity task, we assessed whether the original channels contributing to the PCs varied in amplitude depending on whether the magnitude of tPC_1_ was positive or negative. We used two-sample *t*-tests (*p*<0.0005) to compare the amplitude of each physiological channel during phases in which the magnitude of the tPC_1_ was positive versus phases in which the magnitude of the tPC_1_ was negative (*t.test* package in R). This analysis revealed that periods in which the magnitude of tPC_1_ was positive (i.e., the baseline periods) were characterized by deeper, slower respiration and slower heart rate while periods in which the magnitude of tPC_1_ was positive (i.e., the emotion trials) were characterized by faster, shallower respiration and faster heart rate (fig. S4).

Fractional occupancy of ANS states following k-means clustering of tPCs in individual participants

A two-way analysis of variance that examined the fractional occupancy scores of individual participants revealed a significant main effect of ANS state, *F*(4,1100)=3.2, *p*<0.05, and a significant ANS state by emotion trial interaction, *F*(16,1100)=7.3, *p*<0.0005, which indicated that participants spent different percentages of time in the ANS states during the emotion trials. As in the group-level results (Fig. 2G), k-means clustering in individual participants found that ANS State 1 was the predominant state for most participants during the awe trial; ANS State 2, during the sadness trial; ANS State 3, during the amusement trial; ANS State 4, during the disgust trial; and, to a lesser degree, ANS State 5, during the nurturant love trial (fig. S7).

Amplitude of physiological channels and ANS states

To understand the physiological profile of each ANS state, we conducted a two-way analysis of variance (with random intercepts for participants) to compare the standardized time series concatenated across the sample of each physiological channel across the five ANS states. This analysis revealed significant main effects of ANS state, *F*(4,168425)=3044.3, *p*<0.0005, and physiological channel, *F*(7,352)=44.6, *p*<0.0005, as well as a significant interaction between ANS state and physiological channel, *F*(28,168425)=1472.0, *p*<0.0005. This analysis indicated that each ANS state was characterized by a unique constellation of physiological activity (fig. S8).

Resting Period

*Comparison of the Principal Components from the Resting Period and the Emotional Reactivity Task*

Given that principal component analyses on two distinct data sets can yield similar components but with a different order or sign of eigenvector loadings, we reordered the PCs from the analysis of the resting period in such way that we minimized the cosine distance (D_c_) between the eigenvector loadings from the PCs derived from the resting period and those from the emotional reactivity task (Fig. 3A). In the principal component analysis of the resting period, PC_2_ corresponded to PC_1_ from the emotional reactivity task (D_c_=0.14); PC_3_, to PC_2_ (D_c_=0.06); PC_6_, to PC_3_ (D_c_=0.33); and PC_1_, to PC_4_ (D_c_=0.31). The inversed eigenvector loadings of PC_4_ in the resting period had the highest similarity to the eigenvector loadings of PC_5_ from the emotional reactivity task (D_c_=0.13). PCs_1-4_ and PC_6_ from the resting period ANS data explained 76% of total variance, with each PC explaining at least 9% of variance. We reordered the resting period PCs to align with those of the emotional reactivity task for use in subsequent analyses.

*Comparison of the Resting Period with the Emotional Reactivity Task*

Pearson’s correlation analyses found high spatial similarity between the vectorized eigenvector loadings derived from the resting period and those from the emotional reactivity task, *r*(40)=0.82, *p*<0.0001, (fig. S9B). These findings suggested ANS activity during the resting period and the emotional reactivity task shared similar functional architectures.

Supplementary Figures and Tables

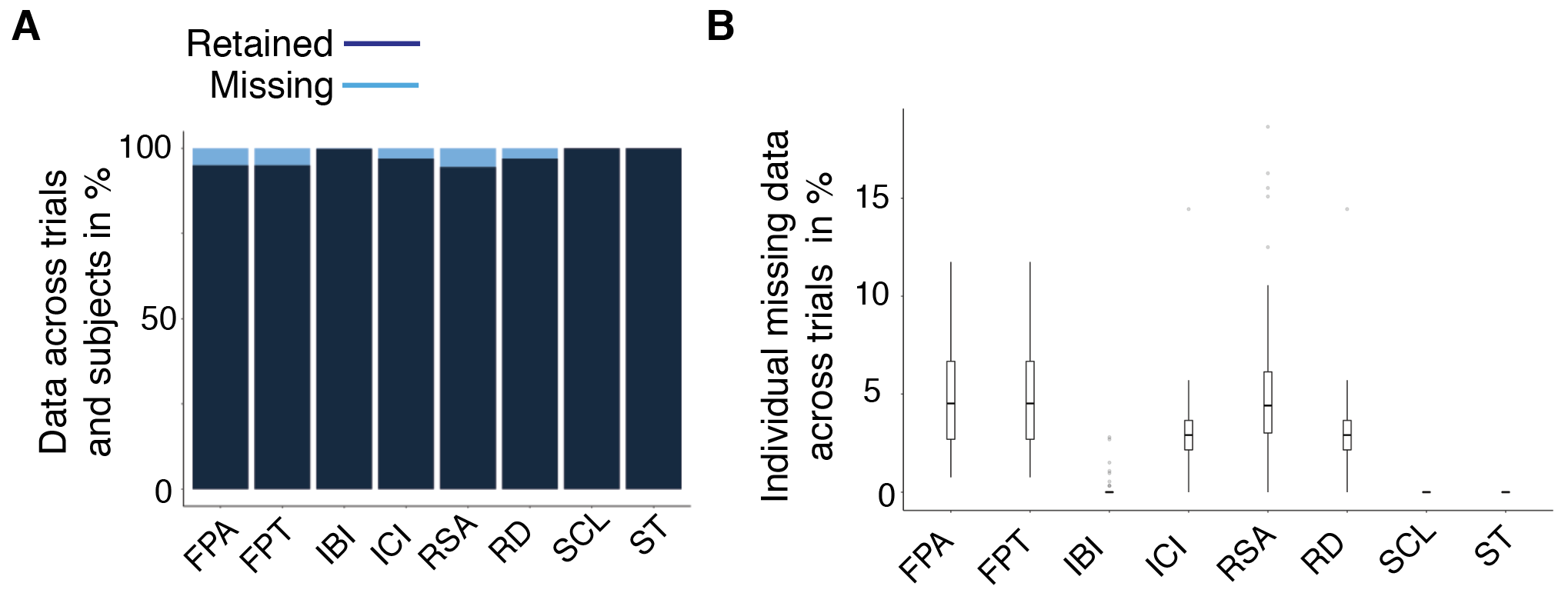

Fig. S1. Interpolation of missing physiological data in the final sample.

(A) In the 45 participants included in the final sample, between 0.2% (IBI) and 5.5% (RSA) of missing physiological data was replaced through spline interpolation. (B) Boxplots show the percentage of missing data points in each physiological channel during the emotional reactivity task; all participants retained at least 80% of the data in any channel. IBI = inter-beat interval; ICI = inter-cycle interval; FPA = finger pulse amplitude; FPT = finger pulse transit time; RD = respiration depth; RSA = respiratory sinus arrhythmia; SCL = skin conductance level; ST = skin temperature.

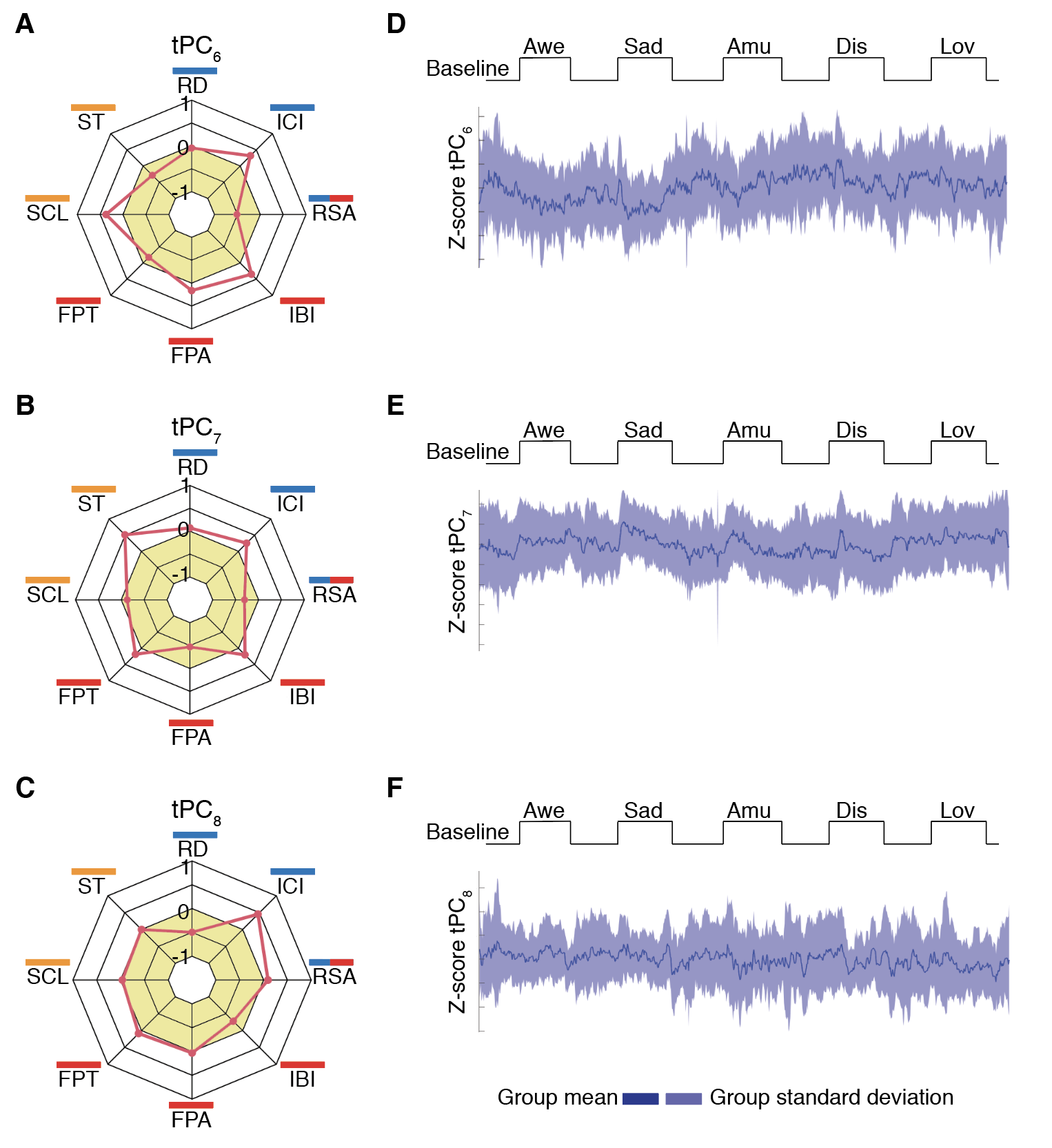

Fig. S2. ANS profiles of principal components six, seven, and eight.

In the ANS data concatenated across the sample, PC_6-8_ each explained less than 10% of variance and were not included in further analyses. On the left, eigenvector loadings are shown for **(A)** PC_6_, **(B)** PC_7_, and **(C)** PC_8_. Radians in ochre represent negative loadings. On the right, the time series of each PC (tPC) was computed and plotted to illustrate second-by-second fluctuations during the emotional reactivity task in **(D)** PC_6_, **(E)** PC_7_, and **(F)** PC_8_. Amu = amusement trial; Awe = awe trial; Dis = disgust trial; IBI = inter-beat interval; ICI = inter-cycle breath interval; FPA = finger pulse amplitude; FPT = finger pulse transit time; Lov = nurturant love trial; RD = respiration depth; RSA = respiratory sinus arrhythmia; Sad = sadness trial; SCL = skin conductance level; ST = skin temperature.

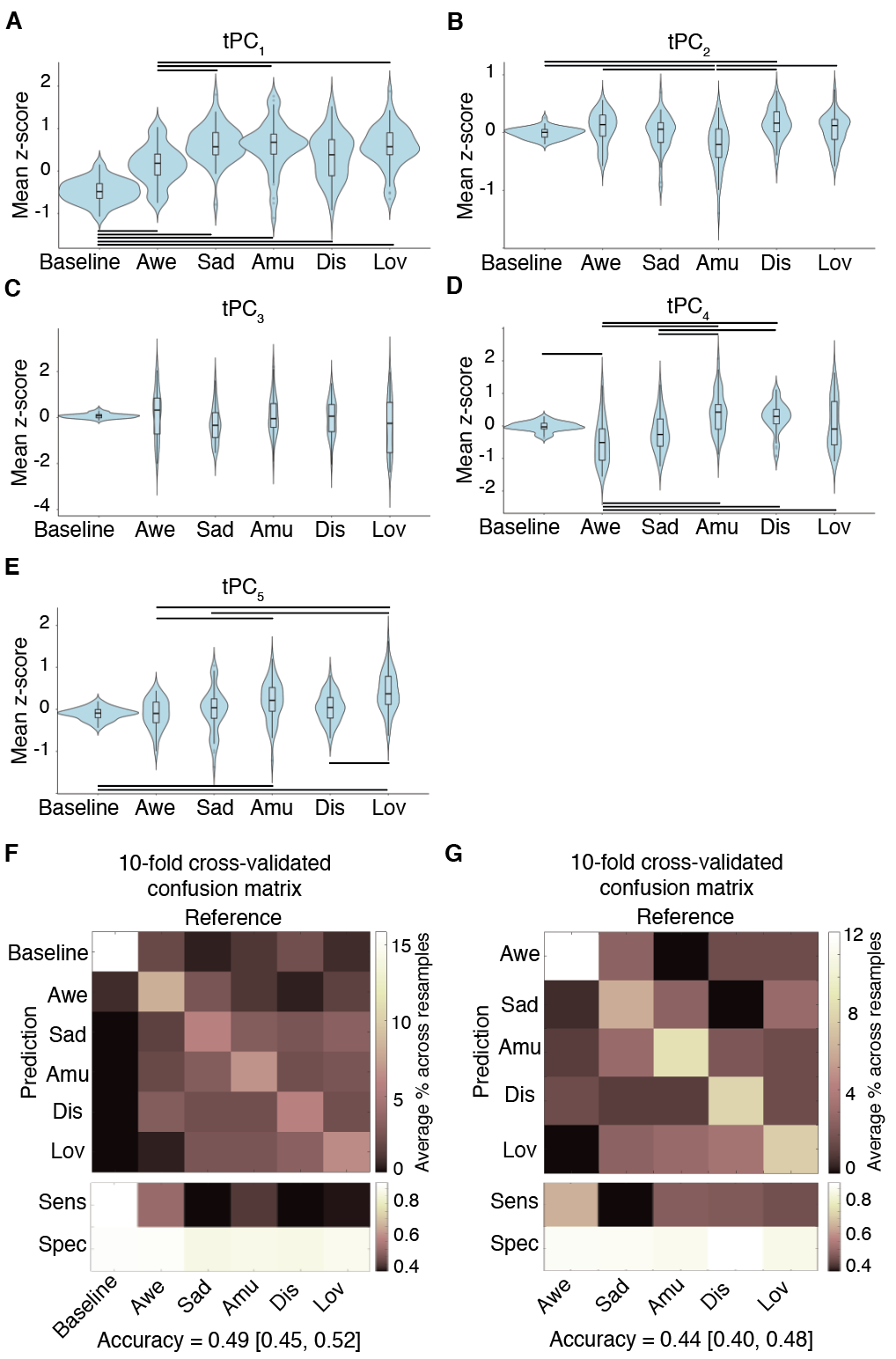

Fig. S3. Mean activity of the principal components during the emotional reactivity task.

(**A-E**) We computed the average magnitude of each tPC during the baseline periods and emotion trials. Each tPC had a different magnitude during the emotion trials. Bars reflect *p*<0.05, Bonferroni-corrected pairwise comparisons. Multinomial logistic regressions were performed to determine whether the averaged tPC magnitudes predicted the trial. Confusion matrices, derived using a 10-fold cross-validation approach, show the correct classification of trials based on averaged tPC magnitudes. Sensitivity (sens), specificity (spec), and accuracy levels [confidence interval in square brackets] are provided for the logistic regression models that included (**F**) the emotion trials and the averaged pre-trial and post-trial baselines; (**G**) the emotion trials alone.

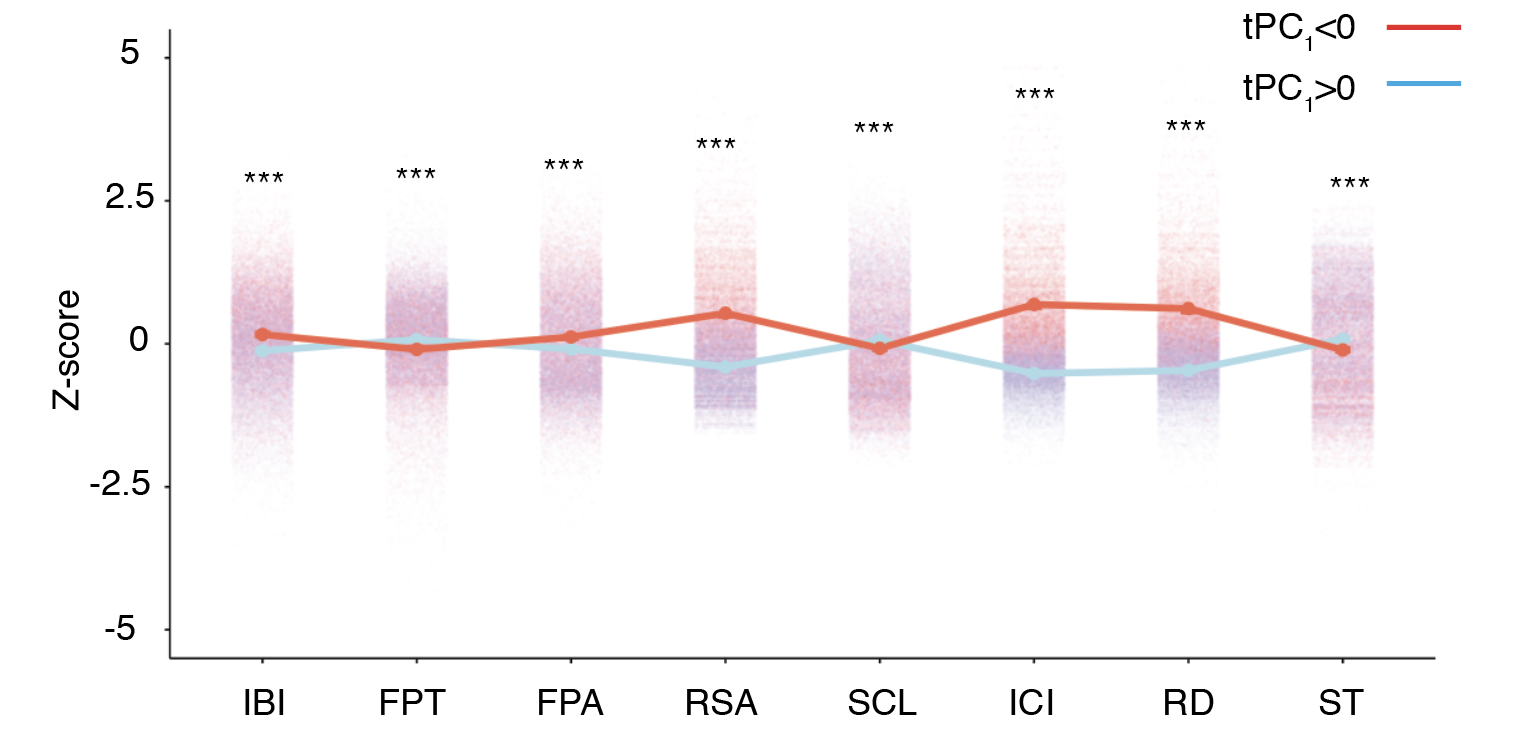

Fig. S4. Activity in individual physiological channels during different phases of tPC_1_.

tPC_1_ oscillated across the baseline periods and emotion trials in a manner that closely aligned with the task structure (Fig. 2A). In general, the magnitude of tPC_1_ was less than zero (i.e., negative *z*-score) during the baseline periods and greater than zero (i.e., positive z-score) during the emotion trials. To investigate the activity levels of individual physiological channels during these different phases of tPC_1_, we coded each data point according to whether it was acquired during a second in which the magnitude of tPC_1_ was less than zero (red line) or greater than zero (blue line). We then used two-sample *t*-tests to compare the amplitudes in each physiological channel that were acquired during these two phases of tPC_1_ (*** indicates significance at *p*<0.0005). Line-plots (showing means and standard errors) with jittered data points show that periods in which the magnitude of tPC_1_ was less than zero, which typically occurred during the baseline periods, were characterized by higher values for inter-beat interval, finger pulse amplitude, respiratory sinus arrhythmia, inter-cycle interval, and respiration depth and lower values for finger pulse transit time, skin conductance level, and skin temperature—an overall pattern that is consistent with higher parasympathetic nervous system activity at rest. This pattern reversed when we examined activity levels in individual physiological channels during periods in which the magnitude of tPC_1_ was greater than zero, which may reflect greater sympathetic nervous system contributions to certain emotion trials.

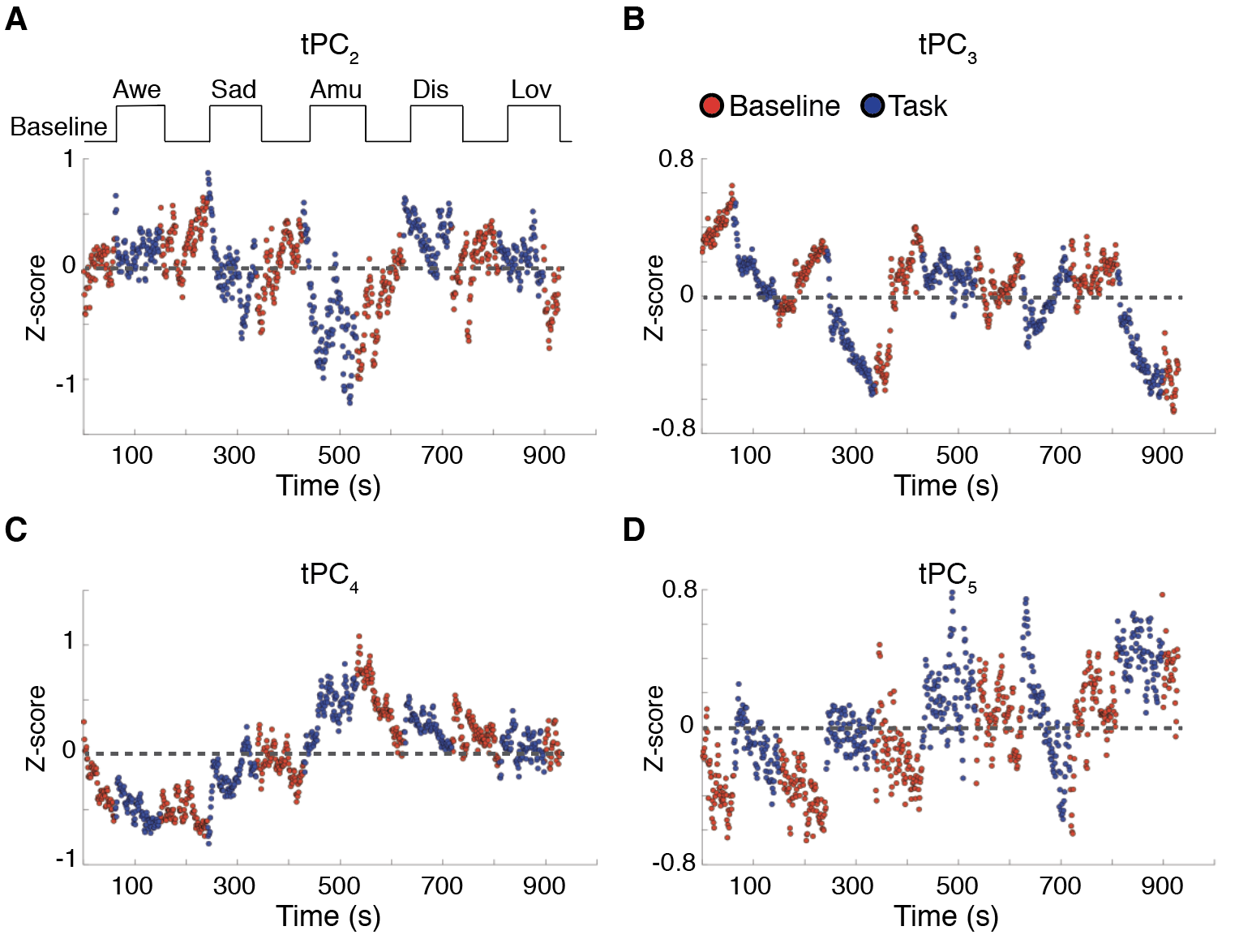

Fig. S5. Amplitude fluctuations of tPC_2-5_ during the baseline periods and emotion trials.

Fluctuations in the time series of principal components 2-5 (tPC_2-5_; A-D) during the emotional reactivity task did not differentiate the baseline periods (“Baseline,” shown in red) from the emotion trials (“Task,” shown in blue) but rather appeared to show trial-specific activity patterns (e.g., during the amusement trial in blue at around 500 sec, tPC_2_ had a negative amplitude and tPC_4_ had a positive amplitude).

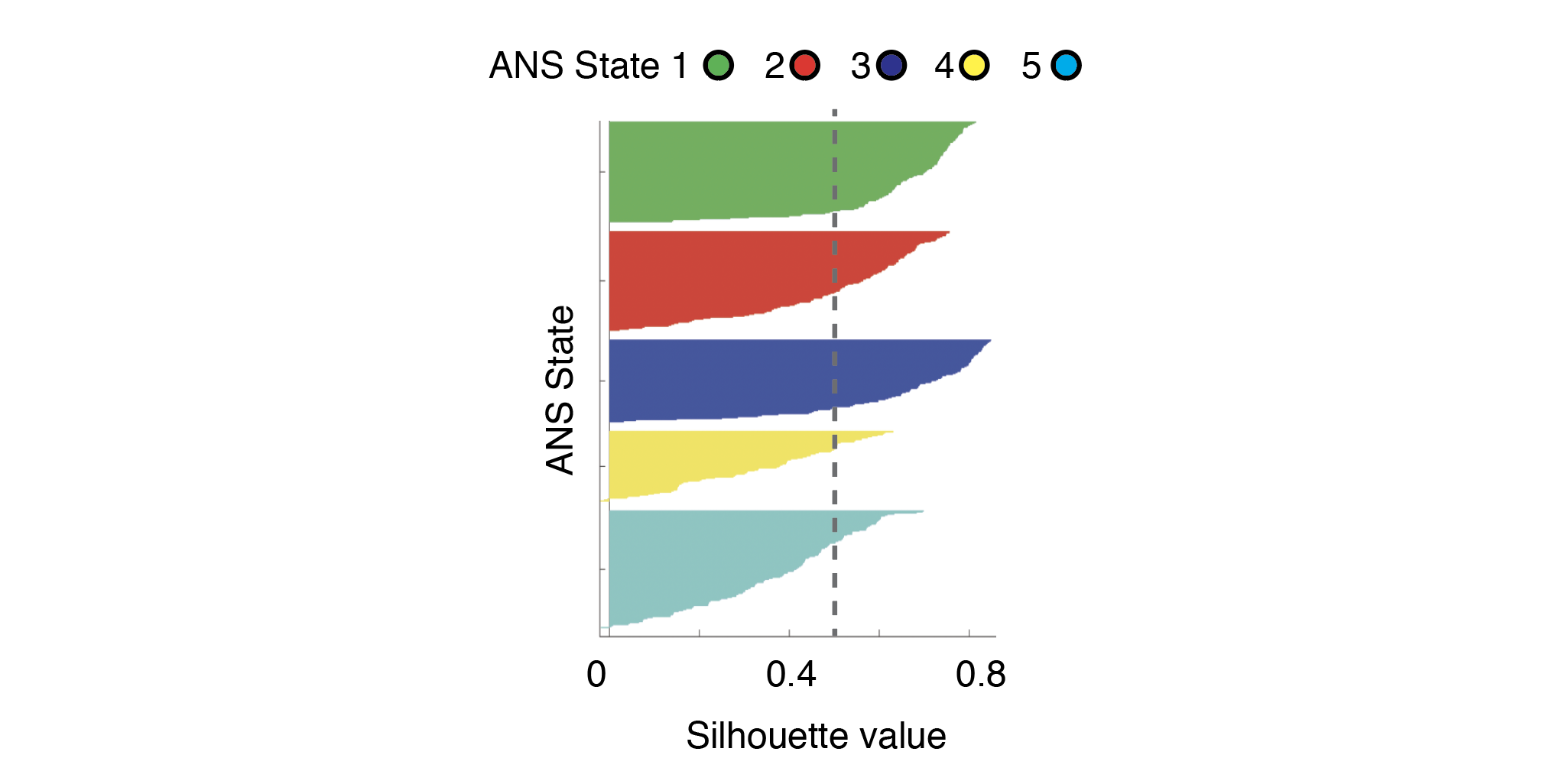

Fig. S6. Silhouette analysis.

A silhouette analysis suggested the presence of five clusters of tPC activity during the emotional reactivity task. The presence of similarly sized clusters (as indicated by the thickness of the silhouette plots) and the fact that all cluster showed above average silhouette values (>0.48, as indicated by the gray dotted line) provided objective support for the k-means clustering partition with k=5 used in our study.

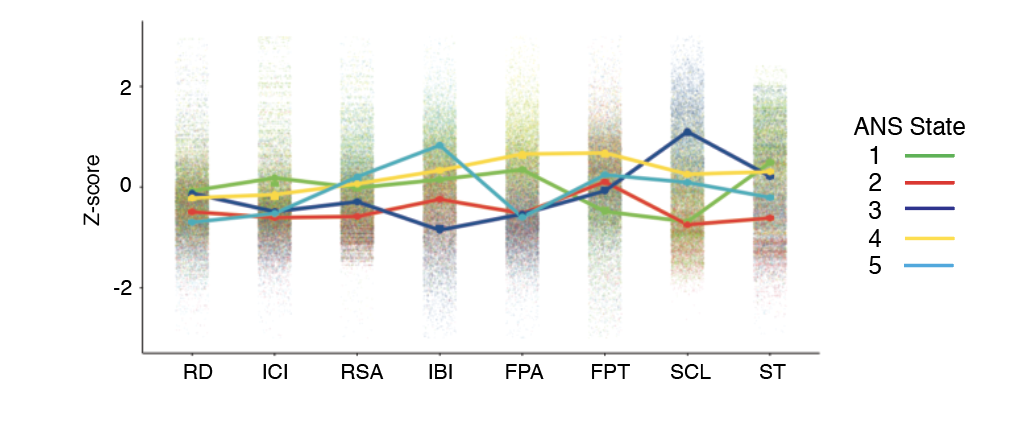

Fig. S7. Amplitude of physiological signals per ANS state.

Line-plots and associated standard error bars display the link between ANS states and the physiological channels that contributed to the original PCs.

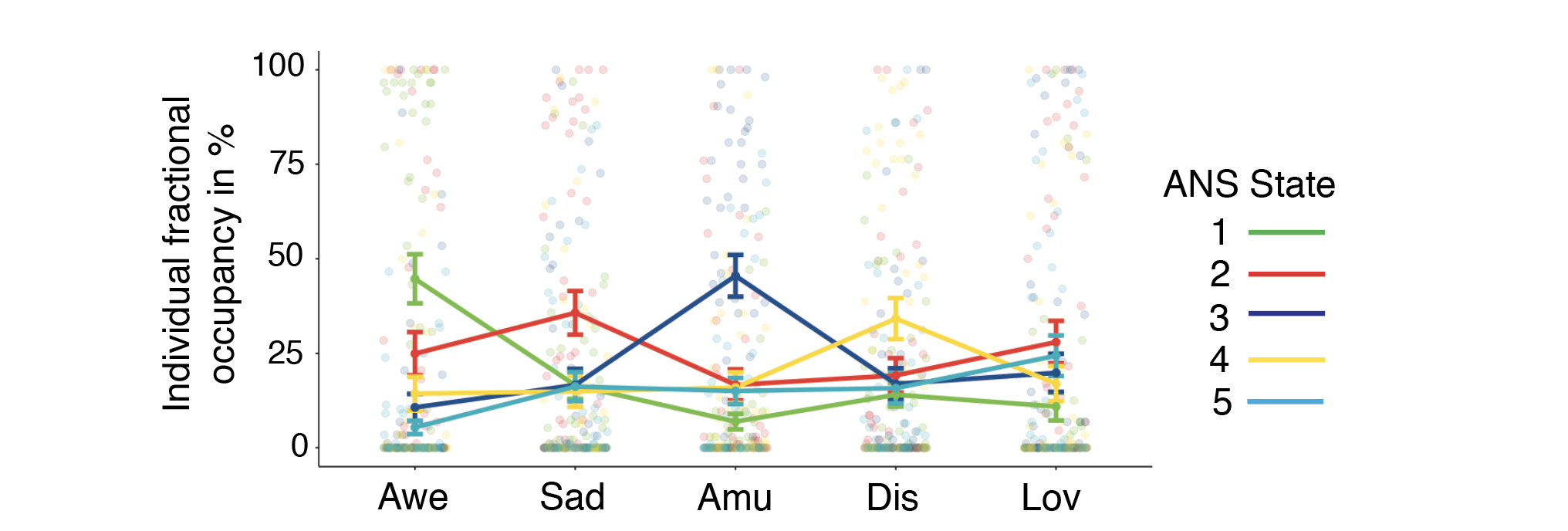

**Fig. S8. ANS state fractional occupancies after k-means clustering of individual participants’ tPCs.**

Each participant’s fractional occupancy scores, the percentage of time spent in each ANS state during each emotion trial, were computed after k-means clustering was performed on the individual tPCs during the emotion trials (awe [Awe], sadness [Sad], amusement [Amu], disgust [Dis], and nurturant love [Lov]). A two-way analysis of variance revealed a significant main effect of ANS state and a significant ANS state by emotion trial interaction on fractional occupancy. Line-plots show the mean fractional occupancy scores of each ANS state across participants during the emotion trials (error bars reflect the standard error). Consistent with the group-level analyses (Fig. 2G), there was a predominant ANS state during each emotion trial when examined at the level of individual participants.

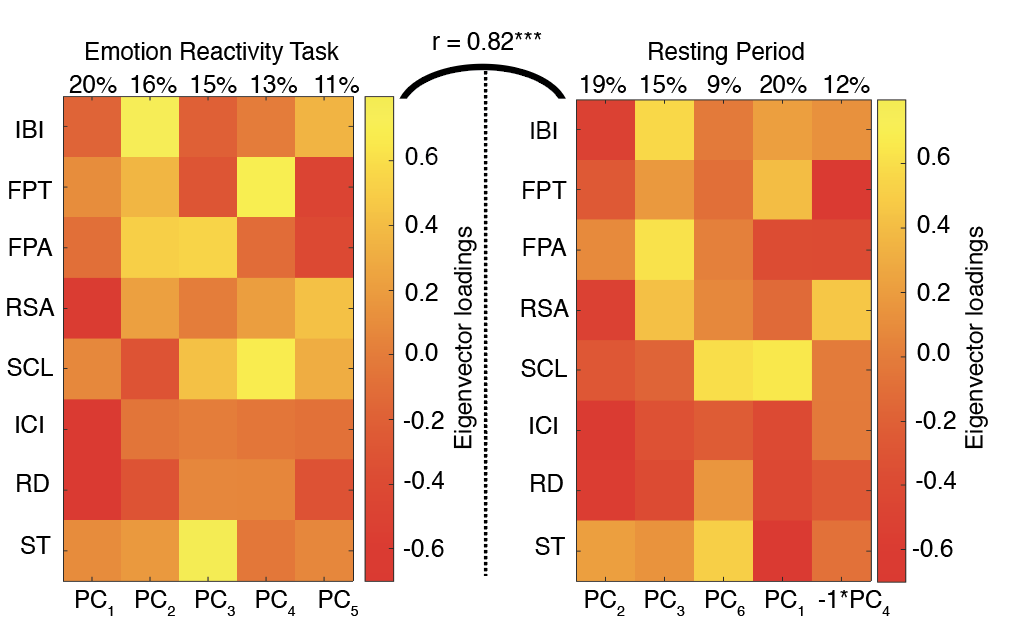

Fig. S9. Correspondence between PC loadings from the emotional reactivity task and the resting period.

Pearson’s correlation analyses found a strong correspondence between the vectorized eigenvector loading matrices derived from the emotional reactivity task (on the left) and the resting period (on the right), *r*(40)=0.82, *p*<0.0001. The values at the top of the heatmaps indicate the percentage of variance explained by each PC.

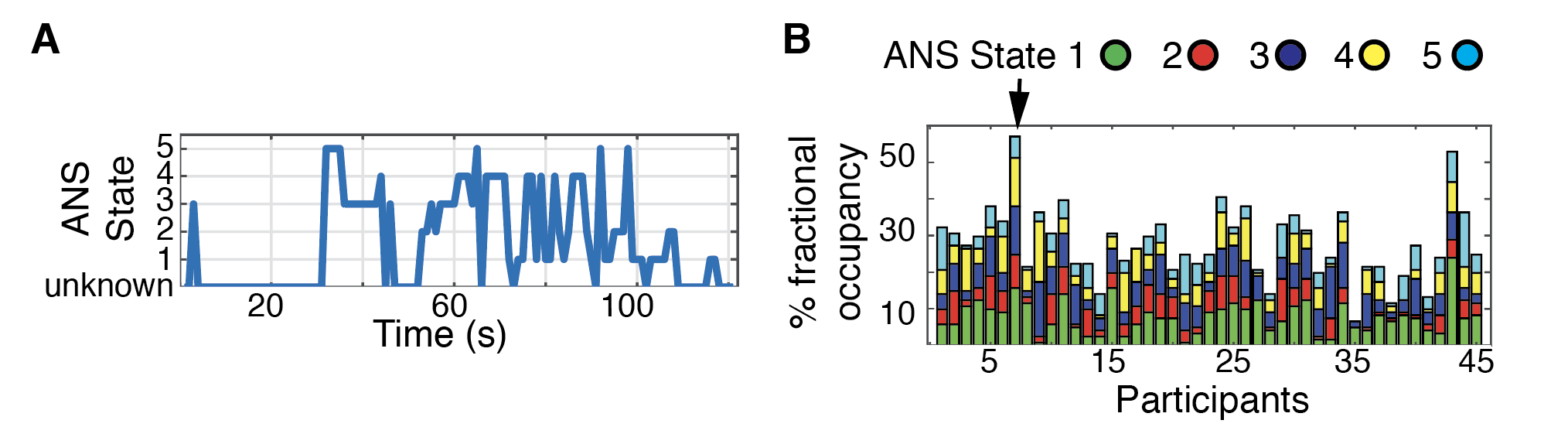

Fig. S10. ANS trajectories of individual participants during the resting period.

(A) The time series of Subject 7’s ANS state occupancy during the resting period is plotted. (B) The stacked ANS state fractional occupancy scores during the resting period are shown for each participant. On average, participants spent 7-57% of the two-minute resting period in one of the ANS states identified during the emotional reactivity task (the arrow points to Subject 7).

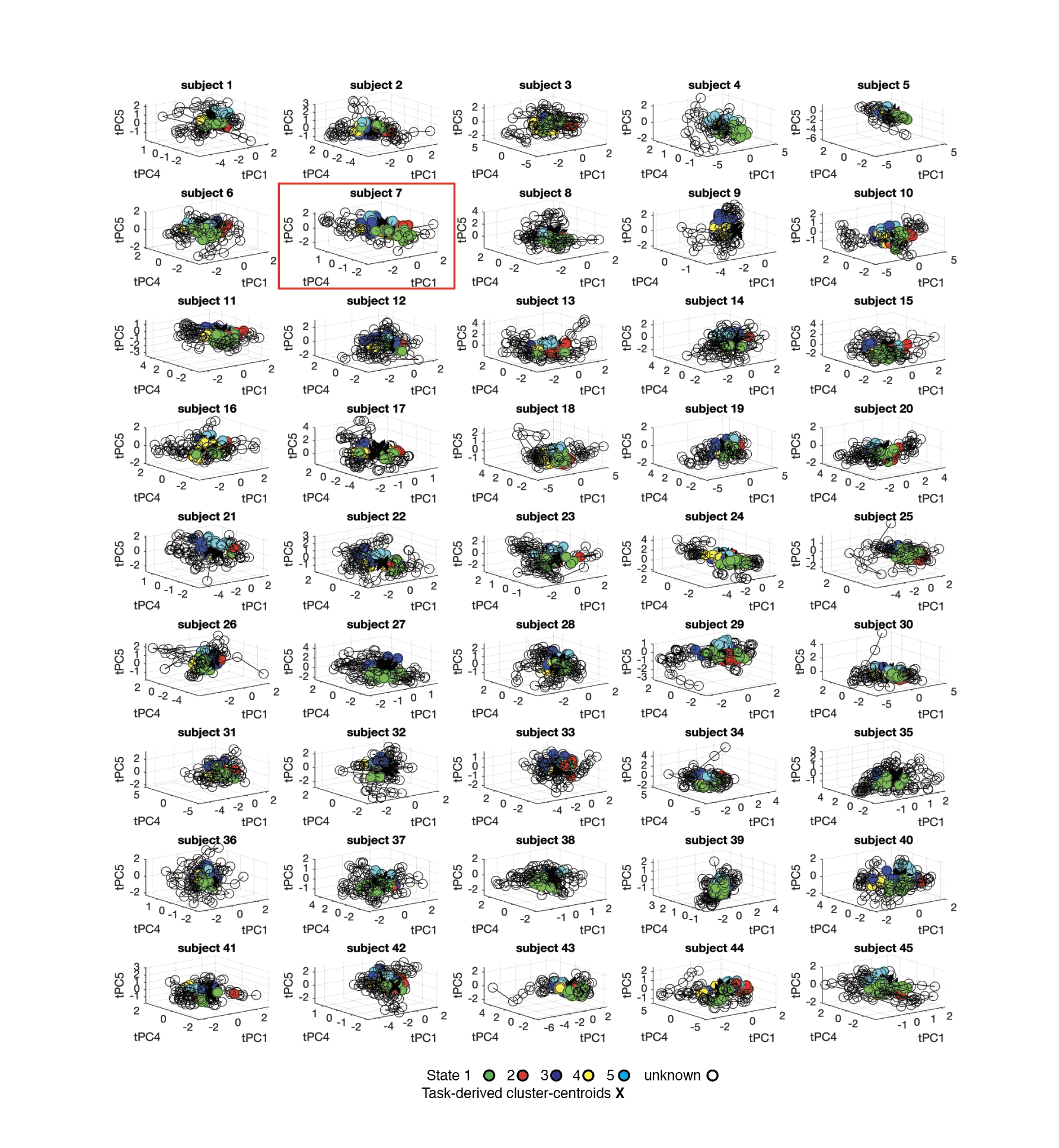

Fig. S11. The ANS trajectories of each participant during the resting period.

For each participant, the trajectories of the tPCs during the resting period are shown here. Data points from the resting period were assigned to ANS states based on the Euclidean distance to the closest cluster-centroids identified in the emotional reactivity task; data points that were not in close proximity to a cluster-centroid were classified as unknown states. Subject 7, highlighted in the main text, is outlined here in red.

|  | *MEAN (SD)* |
| --- | --- |
| SAMPLE SIZE | 45 |
| AGE (YEARS) | 73.6 (4.2) |
| SEX: MALE/FEMALE | 13/32 |
| HANDEDNESS: RIGHT/LEFT | 42/3 |
| EDUCATION (YEARS) | 17.7 (1.8) |
| RACE (ASIAN/ BLACK/WHITE) | 1/1/43 |
| ETHNICITY (HISPANIC OR LATINO) | 1 |
| **MINI-MENTAL STATE EXAMINATION (/30)** | 29.3 (1.0) |
| **CLINICAL DEMENTIA RATE SCALE TOTAL (/3)** | 0.0 (0.0) |
| **BENSON FIGURE 10-MINUTE RECALL (/17)***** | 10.8 (3.4) |
| **MODIFIED TRAILS (CORRECT LINES PER MINUTE) **** | 38.2 (14.4) |
| **MODIFIED TRAILS ERRORS***** | 0.3 (0.7) |
| **PHONEMIC FLUENCY (# CORRECT IN 60 SECONDS) **** | 16.0 (3.9) |
| **SEMANTIC FLUENCY (# CORRECT IN 60 SECONDS)** | 21.8 (3.9) |
| **DESIGN FLUENCY CORRECT (# CORRECT IN 60 SECONDS) **** | 11.6 (4.6) |
| **DIGITS BACKWARD**** | 5.2 (2.1) |
| **BENSON FIGURE COPY (/17)**** | 14.9 (3.3) |
| **BOSTON NAMING TEST SPONTANEOUS CORRECT (/15)**** | 13.7 (4.6) |

Table S1. Demographic and cognitive data for the final sample. For questionnaires, the highest attainable score is shown in the parentheses. SD = standard deviation. *Questionnaire completed by 44 out of 45 participants; **questionnaire completed by 43 out of 45 participants; *** questionnaire completed by 42 out of 45 participants

| *Awe trial* | | | |
| --- | --- | --- | --- |
| Question | Multiple choice answers | | |
| What happened in this movie? | Pictures of forest fires | **Pictures of mountains and oceans** | Pictures of children who were playing |
| % of responses | 0 | 100 | 0 |
| *Sadness trial* | | | |
| Question | Multiple choice answers | | |
| What happened in this movie? | A dog played with a ball | A woman ate dinner | **A woman was crying** |
| % of responses | 0 | 0 | 100 |
| *Amusement trial* | | | |
| Question | Multiple choice answers | | |
| What happened in this movie? | **A baby laughed** | A baby threw a toy | A man blew his nose |
| % of responses | 100 | 0 | 0 |
| *Disgust trial* | | | |
| Question | Multiple choice answers | | |
| What happened in this movie? | A cow ate grass | Someone walked in a park | **Someone cleaned an ear** |
| % of responses | 0 | 0 | 100 |
| *Nurturant love trial* | | | |
| Question | Multiple choice answers | | |
| What happened in this movie? | Cars were driving | A kid went to the doctor | **Babies were playing** |
| % of responses | 0 | 0 | 100 |

Table S2. After each emotion trial, multiple choice questions were presented to assess whether participants played attention to the videos. The correct answer is highlighted in bold. All participants provided the correct response, suggesting that they engaged the emotional reactivity task.

|  |  | Video | | | | |  |  |
| --- | --- | --- | --- | --- | --- | --- | --- | --- |
|  |  | Awe | Sadness | Amusement | Disgust | Nurturant love | *F* | *p*< |
| Emotion self-report, mean (standard deviation) | Affection | 0.6 (0.6) | 0.6 (0.7) | 1.5 (0.6) | 0.1 (0.3) | 1.4 (0.6) | 47.4 | .001 |
|  | Amusement | 1.3 (0.5) | 0.0 (0.0) | 1.9 (0.3) | 0.2 (0.4) | 1.7 (0.5) | 35.2 | .001 |
|  | Anger | 0.0 (0.0) | 0.4 (0.5) | 0.0 (0.0) | 0.0 (0.2) | 0.0 (0.0) | 19.5 | .001 |
|  | Awe | 1.6 (0.5) | 0.1 (0.2) | 0.9 (0.7) | 0.8 (0.8) | 0.7 (0.6) | 37.3 | .001 |
|  | Disgust | 0.0 (0.0) | 0.1 (0.3) | 0.0 (0.0) | 1.1 (0.8) | 0.0 (0.0) | 76.0 | .001 |
|  | Embarrassment | 0.0 (0.0) | 0.0 (0.2) | 0.0 (0.0) | 0.1 (0.3) | 0.0 (0.0) | 4.7 | .005 |
|  | Excitement/Enthusiasm | 0.8 (0.7) | 0.0 (0.3) | 1.0 (0.7) | 0.1 (0.4) | 0.7 (0.7) | 24.5 | .001 |
|  | Fear | 0.0 (0.2) | 0.4 (0.6) | 0.0 (0.2) | 0.4 (0.6) | 0.1 (0.3) | 9.5 | .001 |
|  | Pride | 0.4 (0.6) | 0.0 (0.0) | 1.6 (0.4) | 0.1 (0.2) | 0.2 (0.5) | 5.4 | .001 |
|  | Sadness | 0.3 (0.5) | 1.8 (0.4) | 0.1 (0.4) | 0.1 (0.3) | 0.0 (0.1) | 197.0 | .001 |
|  | Surprise | 0.5 (0.3) | 0.5 (0.6) | 0.7 (0.6) | 1.0 (0.6) | 0.3 (0.5) | 11.9 | . 001 |

Table S3. Self-reported emotional experience during each trial of the emotional reactivity task. The two emotions with the highest intensity ratings in each trial are highlighted in bold and shaded in either deep orange (strongest emotion) or light orange (second strongest emotion).
